# Estimating the rates of crossover and gene conversion from individual genomes

**DOI:** 10.1101/2021.11.09.467857

**Authors:** Derek Setter, Sam Ebdon, Ben Jackson, Konrad Lohse

## Abstract

Recombination can occur either as a result of crossover or gene conversion events. Population genetic methods for inferring the rate of recombination from patterns of linkage disequilibrium generally assume a simple model of recombination that only involves crossover events and ignore gene conversion. However, distinguishing the two processes is not only necessary for a complete description of recombination, but also essential for understanding the evolutionary consequences of inversions and other genomic partitions in which crossover (but not gene conversion) is reduced. We present `heRho`, a simple composite likelihood scheme for co-estimating the rate of crossover and gene conversion from individual diploid genomes. The method is based on analytic results for the distance-dependent probability of heterozygous and homozygous states at two loci. We apply `heRho` to simulations and data from the house mouse *Mus musculus castaneus*, a well studied model. Our analyses show i) that the rates of crossover and gene conversion can be accurately co-estimated at the level of individual chromosomes and ii) that previous estimates of the population scaled rate of recombination 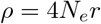 under a pure crossover model are likely biased.

## Introduction

Genetic recombination, the exchange of genetic material between homologous chromosomes during meiosis, is one of the most fundamental evolutionary processes. By creating novel combinations of alleles, recombination increases the efficacy of positive selection (Hill and Robertson, 1966) and reduces the fitness burden of deleterious variants (Charlesworth and Charlesworth, 1997). Recombination breaks down linkage disequilibrium (LD) in the genome and so determines the physical scale over which selective events interfere which each other and affect linked neutral sites (Charlesworth et al., 1993; Simonsen et al., 1996). Since recombination modulates virtually all evolutionary processes, understanding how and why it varies between organisms and between different regions of the genome remains a topic of intense research (see Stapley et al., 2017; Peñalba and Wolf, 2020, for recent reviews). Beyond interest in recombination rate variation *per se*, estimates of recombination are also relevant for other inferences from genomic data. In particular, the power of quantitative or population genetic analyses depends crucially on recombination. Thus, while association studies or inference about past selection (e.g. DeGiorgio et al., 2016, 2014; Setter et al., 2020; Campos and Charlesworth, 2019) and demography (Gutenkunst et al., 2009) often treat single nucleotide polymorphisms (SNP) as independent for the purpose of obtaining point estimates, they rely on parametric bootstrapping or resampling procedures that are conditioned on a model of recombination to quantify uncertainty.

Recombination occurs via double-strand breaks which are either Holliday-junction mediated, resulting in crossovers (CO) and crossover gene conversion (GC) events, or synthesis-dependent strand-annealing mediated, resulting in non-crossover GC events (Resnick, 1976; Szostak et al., 1983; Nassif et al., 1994). In a CO event, two non-sister chromatids break during pairing and reciprocally exchange sequence regions on either side of the break point (Griffiths et al., 2002). In contrast, GC, which typically occurs due to mismatch errors during replication (Carpenter, 1982), involves the non-reciprocal copying of a short stretch of sequence, the GC tract (typically tens to hundreds of bases), from one non-sister chromatid to the other (Szostak et al., 1983; McMahill et al., 2007). The ratio of GC to CO rates varies widely across the tree of life: estimates range from 4-15x in humans (Jeffreys and May, 2004) and mice (Li et al., 2019a) to 1/2 - 1/10x in yeast, algae, and plants (Liu et al., 2018). Similarly, estimates of GC tract lengths range from ten to several thousand base pairs between taxa (Mansai et al., 2011; Casola et al., 2010). Furthermore, the ratio of CO and GC may also vary drastically along the genome. In particular, CO may be severely reduced in centromeric and telomeric regions and within chromosomal inversion, while rates of GC may be unchanged (Talbert and Henikoff, 2010; Korunes and Noor, 2017) or even increased (Crown et al., 2018). However, given that joint estimates for the rates of GC and CO within genomes and across taxa are sparse, the evolutionary causes and consequences of variation in these two components of recombination remain poorly understood.

Much effort has been devoted to estimating CO and GC directly from lab crosses (Hilliker et al., 1994), pedigrees (Kong et al., 2002; Johnston et al., 2016), or sperm-typing (Jeffreys and May, 2004) data. However, such direct estimates are time consuming and expensive given that data from many meiotic events are required. While some pedigree-based (Smeds et al., 2016) and sperm typing methods distinguish CO and GC events, most direct estimates of recombination are necessarily limited to CO events (Kong et al., 2002, 2014; Ma et al., 2015; Johnston et al., 2016). Since individual GC tracts are undetectable unless they span variants, the resolution to detect GC events is inherently limited by the scale of SNP variation.

Given the limitations of direct approaches for estimating recombination, methods that infer recombination indirectly from patterns of LD in whole genome resequence data from natural populations are attractive. LD-based estimators of recombination implemented in popular tools such as *LDhat* (McVean et al., 2002; Auton and McVean, 2007) and *LDhelmet* (Chan et al., 2012) are based on analytic expectations for pairs of loci which, given a large number of pairwise observations, can be used to compute the composite likelihood of the population scaled rate of recombination 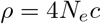. However, current LD based approaches for inferring recombination are limited in at least two ways:

Firstly, both *LDhat* (McVean et al., 2002; Auton and McVean, 2007) and *LDhelmet* (Chan et al., 2012) assume a simple model of recombination that only considers CO and ignores GC. A notable exception is the work of Gay et al. (2007), which extends the copying model of Li and Stephens (2003) to co-estimate CO and GC rates. Secondly, since two-locus approaches are generally conditioned on variant sites, they require phased data from many samples. Such data are still only available for a small minority of taxa.

Here, we address both these limitations by developing a simple composite likelihood method that allows co-estimation of CO and GC rates from individual diploid genomes. The calculation is based on analytic expectations for observing heterozygosity at two loci under the simplest model of recombination (crossover only) and genetic drift (Strobeck and Morgan, 1978; Lohse et al., 2011) and has previously been implemented by Haubold et al. (2010). We first use coalescent simulations to demonstrate that GC biases estimates of the CO rate and show that this bias depends on the physical distance between loci. We then exploit this non-linear dependence of GC on distance by incorporating GC into the two locus expectations which gives a framework for co-estimating the rates of CO and GC. Finally, we apply our method to genome-wide data from wild-caught individuals of the house mouse *Mus musculus castanaeus* and compare our estimates to previous estimates of recombination based on a CO only model (Booker et al., 2017). We also quantify the power of our approach using simulations and investigate the extent to which the rates of CO and GC are correlated with each other and with chromosome length.

## Materials and Methods

### Analytic expectation of two locus heterozygosity

We extended the models of Strobeck and Morgan (1978) and Haubold et al. (2010) to account for GC and use a composite-likelihood approach to co-estimate the rates of CO and GC and the mean GC tract length from individual genomes. We consider a neutral Wright-Fisher model for the evolution of two linked loci separated by *d* nucleotides in a population of *N* diploid individuals. Mutations occur at per-base rate *μ*. CO occurs at per-base rate *c* and results in an exchange of genetic material between sister chromatids. GC initiates at per-base rate *g* and GC tracts are replaced by the sequence from the sister chromatid. For the analysis, we re-scale time by 1*/*2*N* generations and use the population-scaled parameters *θ* = 4*Nμ*, *κ* = 4*Nc*, and *γ* = 4*Ng* for mutation, CO, and GC rates, respectively. We follow Wiuf (2000) in assuming that the GC tract length is an exponentially distributed random variable with mean *L* base pairs (Hilliker et al., 1994; Wiuf and Hein, 2000).

The heterozygosity at a single site *H* with 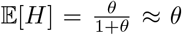 is informative only about the depth of a local genealogy: a site is more likely to be heterozygous when the time to the most recent common ancestor, *T*_*mrca*_, is large and homozygous (i.e. identical in state) when *T*_*mrca*_ is small. Consider a second site at a fixed distance *d* and define *H*_0_, *H*_1_, and *H*_2_ as the proportion of all such pairs where neither site, one site, or both sites are heterozygous (respectively). These two-locus measures of heterozygosity are informative about the joint distribution of the two underlying genealogies and allow estimation of the rate of recombination (Haubold et al., 2010; Lohse et al., 2011).

Using Eq. 4 of Strobeck and Morgan (1978), Haubold et al. (2010) derive analytic expressions for the expected frequency of *H*_0_, *H*_1_, and *H*_2_ as a function of *ρ*, the total rate of events which lead to recombination between two sites separated by *d* base pairs, agnostic to the underlying contributions of CO and GC:

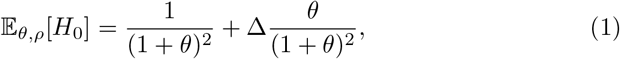

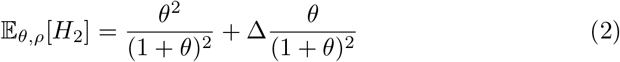

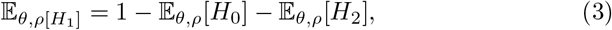

where

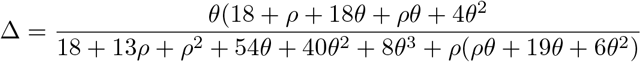

represents the *zygosity* correlation (Lynch, 2008) or the deviation from independence due to linkage. For large *d*, the genealogies at the two sites become independent, so 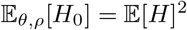 and 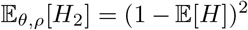, the first term in equations 1 and 2. In contrast, if the second site is tightly linked, the two sites likely share the same genealogy and we expect to see an excess in *H*_0_ and *H*_2_. As *d* increases, so too does the probability that recombination occurs between the sites, resulting in differing genealogies and an increase in *H*_1_.

### Co-estimating crossover and gene conversion rates

The above expectations for *H*_0_, *H*_1_, and *H*_2_ make no assumption about the nature of recombination between pairs of sites, and *ρ* represents the rate at which the alleles transfers to different genetic backgrounds. For sites separated by a given distance *d*, we can obtain a maximum likelihood estimate for the total rate of recombination observed over these distances. Let *n*_*d*,0_, *n*_*d*,1_, and *n*_*d*,2_ be the counts of pairs in the genome corresponding to *H*_0_, *H*_1_, and *H*_2_ for a given distance *d*. The log-likelihood is then

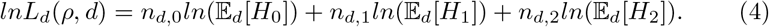

If we assume that recombination between the two loci occurs only through CO, recombination always transfers alleles onto different genetic backgrounds, and the per-base recombination rate *ρ*/*bp* is constant. This is not true for GC, because the two sites will still share a genealogy if the GC tract both initiates and terminates between them. In other words, recombination through GC occurs only if the GC tract spans only one of the two focal sites, in which case GC has the same effect as a CO event. Accounting for the probability of recombination during GC (Wiuf and Hein, 2000; Wiuf, 2000), we can re-write the total rate of recombination *ρ* as a function of distinct rates of CO (*κ*) and GC (*γ*) and the expected GC tract length *L*(Frisse et al., 2001; Langley et al., 2000).

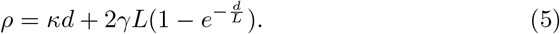

Given the dependence on distance *d*, observations *n*_*d*,0_, *n*_*d*,1_, and *n*_*d*,2_ for a single *d* are insufficient to estimate a three-parameter model of recombination. However, by compositing the likelihood over many distances and substituting Eq. 5 into Eq.s 1 – 3, we can co-estimate the rate of CO *κ*, the rate of GC *γ*, and the mean tract length *L*.

The composite likelihood is thus given by

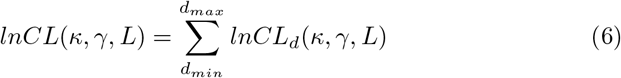

We have implemented the composite likelihood estimation described above in python as a simple open source tool, `heRho` which is available at https://github.com/samebdon/heRho.

### Estimating recombination rates in the eastern house mouse

As a proof of principle, we tested our composite likelihood estimation of recombination on whole genome data from a well studied model species, the eastern house mouse *M*. *m*. *castaneus*. Both direct and indirect estimates for the total rate of recombination exist for this species (Booker et al., 2017) and several studies provide estimates for GC tract lengths (Paigen et al., 2008; Mansai et al., 2011; Li et al., 2019b; Cole et al., 2014).

The data – originally described in Halligan et al. (2010) (ENA accession number PRJEB2176) – consists of Illumina (PE) resequence data for ten individuals sampled from a wild *M*. *m*. *castaneus* population in India. Variant calling is described in (Booker et al., 2017).

To minimize potential biases arising from background selection and the effect of selection on nearby linked sites, all analyses were restricted to intronic regions, which are putatively neutral. Specifically, we considered all introns ¿1kb. Given that the X chromosome has a different recombination rate than the autosomes due to its lack of recombination in males, it was excluded from this analysis.

The final dataset included a total of 123,488 introns on autosomes 1-19, spanning a total of 9 × 10^8^ bases. For each intron, the positions of heterozygous sites in each individual were converted into two-locus counts *n*_*d*,0_, *n*_*d*,1_, and *n*_*d*,2_ for each distance *d* using a custom *Python* script (available on github). `heRho` obtains maximum composite likelihood (MCL) estimates for *ρ* were obtained using the *Python* library*NLopt*.

### Power analysis

To quantify how accurately CO and GC rates can be estimated, we performed a power analyses and parametric bootstrap on data simulated under the full model in `msprime` (Kelleher et al., 2018): we simulated 100 replicates for each chromosome under the MCL estimates obtained from the house mouse data (see Results). Each replicate consisted of 10 diploid samples assuming *θ* = 0.071 (the observed heterozygosity), *μ* = 5.410^−9^ (Uchimura et al., 2015), *L* = 108.4 for all chromosomes. The rates of CO and GC were set to those inferred for each *M*. *m*. *castaneus* chromosome and the length of simulated sequence corresponded to the total length of intronic sequence analysed for each chromosome (simulation code is available in the github repository).

## Results

### Gene conversion explains the non-linear relationship between estimates of *ρ* and physical distance

As a first step, we used Eqs 1 – 4 to investigate how the per-base rate of recombination between pairs of sites depends on the distance between them (Fig. 1). Given a model that only includes CO events (red, dashed), we expect estimates of *ρ*/*bp* to be constant with respect to the distance between sites (red, dashed). However, when GC is included, nearby sites experience a higher per-base rate of recombination than pairs that are distant or unlinked (blue, dashed).

**Figure 1:**
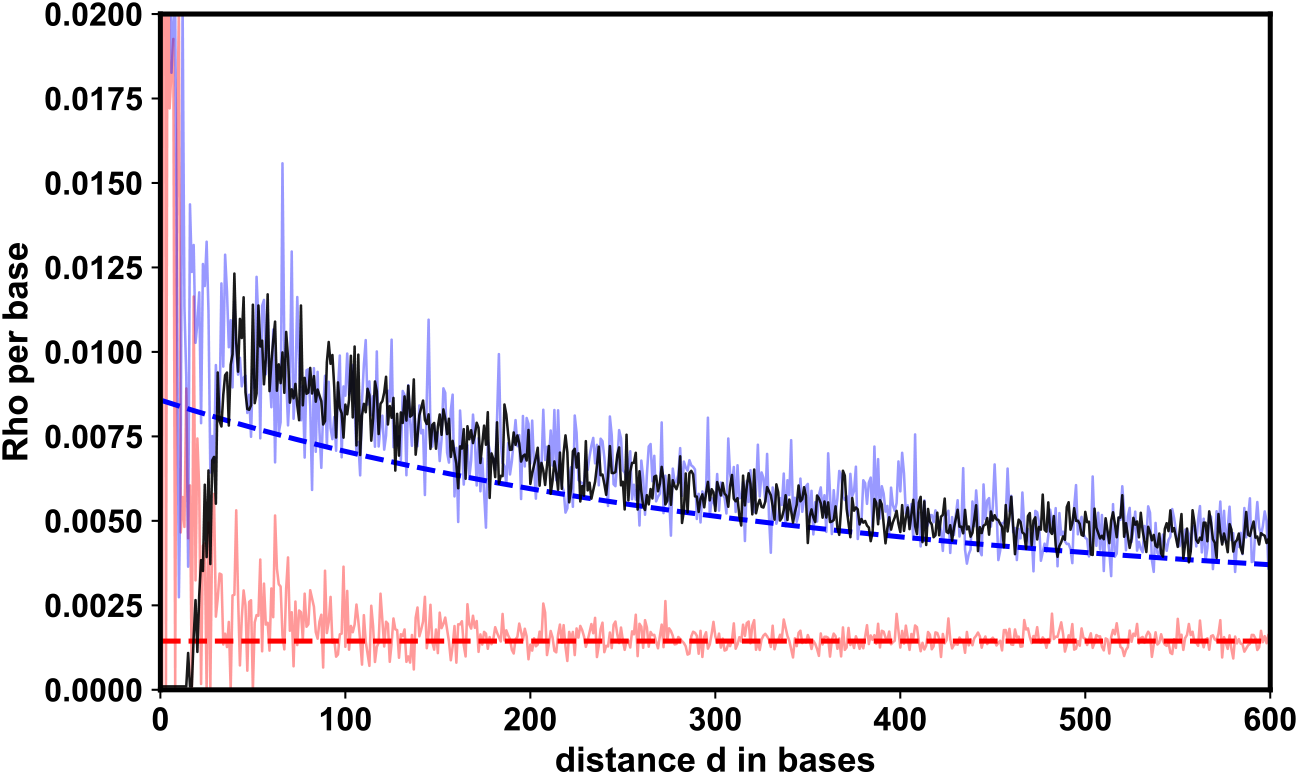
Maximum composite likelihood estimates of *ρ*/*bp* at fixed distances *d* between pairs of sites; simulations with CO only (red), simulations with CO and GC (blue), and empirical data for *M*. *m*. *castanaeus* chromosome 19 (black). In each case, data was combined across a sample of 10 individuals. For simulations, *θ* = 0.0071, *κ* = 0.0014, *γ* = 0.0036, and *L* = 200. The dashed lines show the expectation under the corresponding model (Eq.’s 1 – 4).

We find that estimates of *ρ*/*bp* based on a single replicate simulation either under a model with GC (solid blue) or without (solid red) follow the expected relationship with the distance between sites *d*. When inferring *ρ*/*bp* between pairs of loci at different distances *d* in the mouse data (Fig. 1, black line), the relationship between *ρ*/*bp* and *d* is similar to that seen for data simulated under a model of recombination that includes both CO and GC. We note that the pattern of distance dependent *ρ*/*bp* is not exclusive to *M*. *m*. *castanaeus*, but has been inferred previously by Haubold et al. (2010) for the ascidian *Cionia intestinalis* (although the authors do not mention GC).

Comparing the distance profiles of *ρ*/*bp* estimates between simulated and real data to each other and to analytic expectations (Fig. 1), we find two striking patterns:

Firstly, in the real data, estimates of *ρ*/*bp* are close to zero for nearby pairs of sites and increase sharply over the first ≈ 50 bases. In contrast, while the accuracy and precision of *ρ*/*bp* estimates in simulated data is strongly dependent on *d* (i.e. there is high variability for estimates over short distances *d* < 50), we find no similar monotonic increase in *ρ*/*bp* estimates over the first *approx*50 bases. This discrepancy in estimates of *ρ*/*bp* in real vs simulated data suggests that over short distances *ρ*/*bp* estimates in the real data are biased downwards due to data quality/filtering effects: tightly-linked polymorphisms are difficult to distinguish from complex mutations (e.g. indels) and/or are removed by so called “best practices” variant calling/filtering approaches, skewing the observed values of *H*_0_, *H*_1_, and *H*_2_. This is compatible with the findings of Haubold et al. (2010) who have layered a sequencing error profile on coalescent simulations with CO only and shown that the noise generated by low-coverage data leads to a downward bias in estimates of *ρ*/*bp* over short distances.

Secondly, we find that estimates of of *ρ*/*bp* are generally upwardly biased compared to expectations (compare solid and dashed lines in Fig. 1) due to simplifying assumptions about the mutational process: unlike real genomes which consist of a discrete number of bases, equations 1–3 assume a continuous genome that evolves under the infinite sites mutation model. In that case, the occurrence of two mutations at a pair of sites always results in an *H*_2_ state. In contrast, under a finite-sites mutation model (which `msprime` assumes by default) a back-mutation could generate an *H*_0_ state. Indeed, the resulting upward bias is observed only in simulations that assume finite sites (Fig. S1). The bias is strongest at short distances, where recombination is rare and the expected values of the *H*_*i*_ are primarily governed by the mutational process. However, at greater distances, recombination primarily drives *H*_*i*_ counts and estimates converge to the model predictions.

### Co-estimating crossover and gene conversion rates

By decomposing recombination to distinguish CO and the distance-dependent effects of GC (Eq. 5) and compositing the likelihood over the single-distance counts of *H*_*i*_ (Eq. 6), we may co-estimate both the rates of CO (*κ*) and GC (*γ*) and the mean tract length *L*. However, there are two challenges in implementing this inference: i) the noisiness of the data and the inaccuracy of the analytic results at short distances and ii) the inherent difficulty of co-estimating strongly correlated parameters.

Given the biases over short-distance in the real data, an obvious strategy is to introduce a minimum distance *d*_*min*_ in the composite likelihood (Eq. 6). However, since most of the information to co-estimate the rate of GC and the mean GC tract length is contained in short-distances, there is a trade-off between minimizing bias and retaining information. Our exploration of this trade-off both in real and simulated data shows that parameter estimates are stable across a broad range of *d*_*min*_ values (Fig. S2). To minimize the loss of information, we chose *d*_*min*_ = 100*bp* for all further analyses. Since genomes are finite and analysis is often restricted to a particular genomic partition, an upper distance threshold *d*_*max*_ is also unavoidable. To avoid biasing inference towards very long introns (which are selectively constrained), we limited the analysis to the first *d*_*max*_ = 1000*bp* of each intron.

For our chosen range of distance thresholds (*d*_*min*_, *d*_*max*_) = (100, 1000), as the next step in our preliminary analysis we examined the of the mouse data, we asked whether sufficient information is retained to confidently co-estimate the three recombination parameters. To do this, we focused on chromosome 19 for which `heRho` gives the following maximum composite likelihood estimates: *κ* = 0.00267, *γ* = 0.0044, and *L* = 113.24. Examining the support, as measured by the logarithm of the composite likelihood (*lnCL*), surface around this maximum illustrates the challenge of co-estimating *L* and *κ*. Although estimates are negatively correlated (Fig. 2 B), we were positively surprised that it is not only possible to co-estimate both parameters (the *lnCL* surface is smooth and contains a distinguishable optimum, Fig. 2), but that the estimates are indeed plausible, i.e. are compatible with direct, experimental estimates.

**Figure 2:**
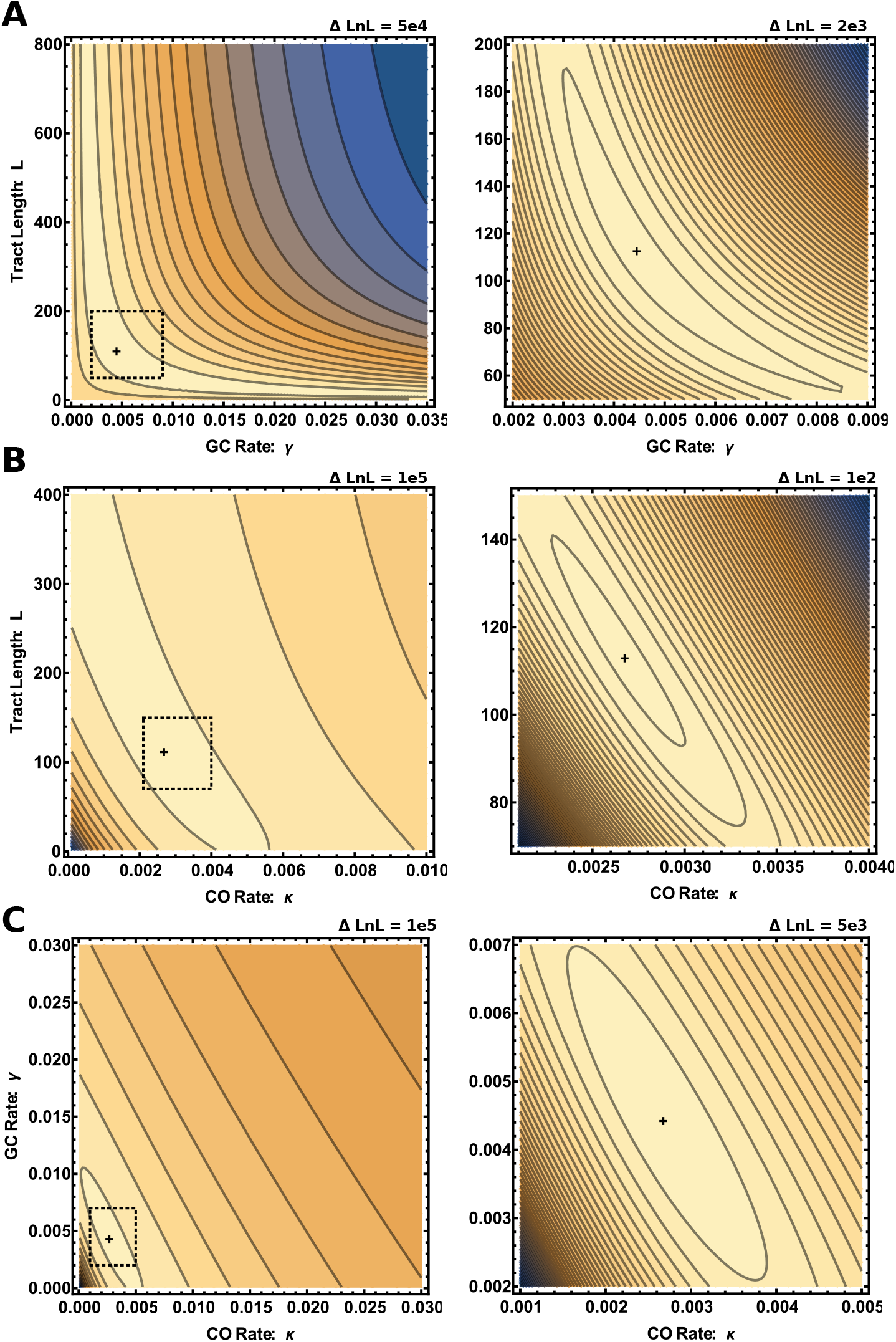
The composite likelihood surface for the rates of CO, GC, and the mean length of GC tracts for intronic data from *M*. *m*. *castaneus* chromosome 19. Each panel shows the two-dimensional projection of the composite likelihood surface (*lnCL* increases from blue to yellow) under the global MCL estimates of parameters: 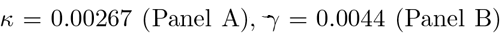, and *L* = 113.24 (Panel C). For each panel, a broad parameter region is shown in the left plot, while the right plot focuses on the region near the optimum indicated by the dashed square. In all plots the distance between contours is indicated at the top and the cross denotes the MCL estimate.

Less surprisingly, we observe that estimates for the GC rate *γ* and the mean tract length *L* are negatively correlated both with each other (Figure 2 A) and with estimates of the CO rate *kappa* (Figure 2 B and C). Given the degree to which parameter estimates are confounded and the fact that we have no biological reason to expect the length of GC tracts to vary between chromosomes, we chose to co-estimate a global *L* and chromosome-specific GC and CO parameters in the subsequent analysis of the mouse data described below.

### The recombination profile of *M*. *m*. *castanaeus*

Our per chromosome co-estimates of the rates of CO (*κ*) and GC (*γ*) in *M*. *m*. *castaneus* (based on data from all 10 individuals, Fig. 3) range from 0.00145 to 0.00269 and 0.00211 to 0.00461 and respectively. Assuming that the mean GC tract length is the same for all autosomes, our global MCL estimate for this parameter is 108 bases, which is within the range of previous direct estimates (≈ 10-300 for NCO and 200-1200 for CO gene conversion tracts (Paigen et al., 2008; Mansai et al., 2011; Li et al., 2019b; Cole et al., 2014)). When restricting our analysis to a single individual, we obtain broadly concordant estimates of the CO rate and mean tract length, but with less data available, the GC rate estimates vary much more across chromosomes (Fig. S3). Indeed, for simulations, increasing the sample size from one (Fig. S4) to ten individuals (Fig. 3) reduces the variance but not the bias of the results.

**Figure 3:**
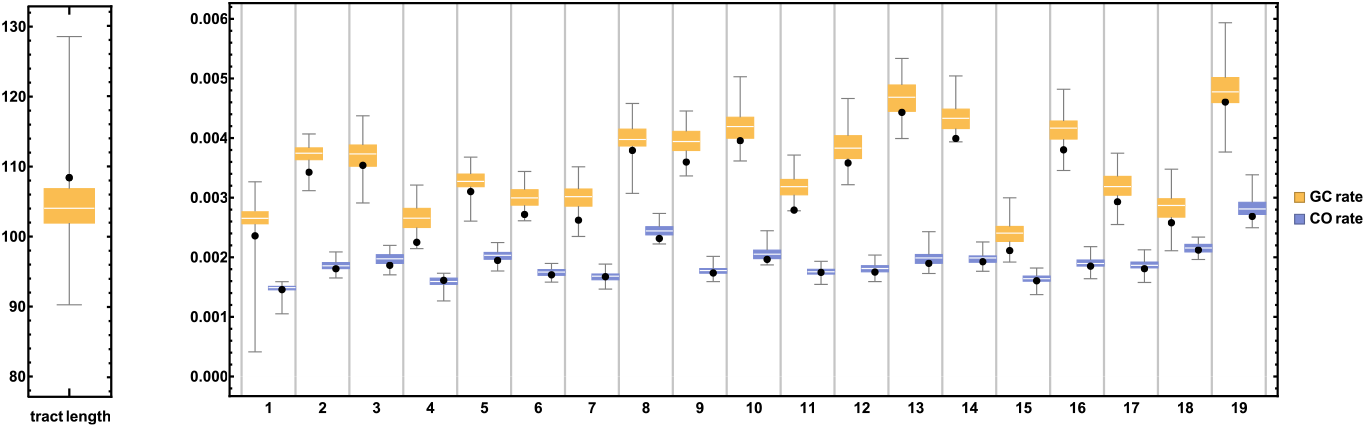
Recombination parameters co-estimated for the 19 autosomes of *M*. *m*. *castanaeus* using data pooled across ten individuals (black dots) and corresponding parametric bootstrap results from 100 replicate simulations. The per chromosome estimates of the GC rate (*γ*) and mean tract length are shown in yellow, estimates for the rate of CO (*κ*) in blue.

Our per chromosome estimates recover several well known, broad-scale patterns: Firstly, as some GC events occur during CO, we expect the rates of CO and GC to be mechanistically and positively correlated, and this is indeed the case (Figure S5). Note that this signal contrasts with the negative correlation in the estimation error of both parameters (Fig. 2 C) and therefore must reflect the underlying dynamics of meiotic recombination rather than any statistical artefact.

Secondly, as chromosomes have a minimum bound of map length at 50cM due to obligate CO, we expect the CO rate per-base to be negatively correlated with chromosome length. We recover this pattern (Figure S5) that is widely documented not only in mammals (Johnston et al., 2017), including humans (The International Genome Sequencing Consortium, 2001), but also in flycatchers (Kawakami et al., 2014), yeast (Kaback et al., 1999) and butterflies (Martin et al., 2016). In contrast, we find that per chromosome estimates of the rate of GC are not significantly correlated with chromosome length (*p* = 0.148) (Figure S5 C). Since a high proportion of GC products are the result of non-CO recombination events, we do not expect GC rates to correlate significantly with patterns of chiasma formation.

## Discussion

A significant challenge in population genetics is to develop inference methods that are both efficient in extracting signals about population processes from sequence variation and simple, i.e. rely on a minimum number of assumptions. Given that high-coverage whole-genome data have become the norm, we now have the ability to study the fundamental forces of evolution, such as recombination, both at fine genomic scales and across a broad taxonomic range. We have developed a method for quantifying CO and GC from the distribution of heterozygous sites in small samples – even from individual diploid genomes.

### heRho’s strengths and weaknesses

Not only does our framework allow for more complete/realistic estimates of recombination than previous CO-restricted methods (Auton and McVean, 2007; Haubold et al., 2010), it is also simpler and less error prone since it does not rely on phase information which can bias results. For example, Booker et al. (2017) find that in the presence of switch errors, LDhelmet consistently overestimates the CO rate. Furthermore, by including homozygous states in the analysis, we garner sufficient information to co-estimate CO and GC when data are restricted to short distances. As such, `heRho` can potentially generate a whole-genome annotation-specific recombination profile, even for small genomic features that partition the genome (e.g. 4D sites in exons).

Our method does suffer from many of the same potential biases as other population genetic estimators of recombination: given that we are assuming a neutrally evolving Wright-Fisher population of constant size, any demographic and selective processes that generate LD (e.g. positive selection, population bottlenecks or admixture), will bias estimates of recombination obtained with `heRho` downwards. Perhaps a more important question is whether there are processes that bias the inference of GC rates specifically or, in the extreme case, may mimic signatures of GC, where none exist. While there are processes that may increase short-range measures of LD (e.g. heterogeneity in mutation rate that leads to an excess of *H*_2_ and *H*_0_), we cannot think of forces or mechanisms that, like GC, have the opposite effect of decreasing LD only over short genomic distances on the order of GC tract lengths.

As demonstrated, our method `heRho` relies on large amounts of sequence data and is fundamentally limited by the frequency of the rarest two-locus observation *H*_2_, which for any distance *d*, is of order *H*^2^. We therefore expect that it will not be possible to obtain estimates of GC and CO at small genomic scales (say in windows of 100kb). While pooling observations across individuals increases the number of *H*_*i*_ observations and reduces variance in the estimates, we expect many heterozygous sites to be shared among individuals, and thus the returns diminish quickly with sample size.

### Reconciling heRho’s recombination estimates for *M*. *m*. *castaneus* with LDHelmet

How do our estimates in *M*. *m*. *castaneus* compared to those obtained previously by Booker et al. (2017) using LDHelmet and a CO-only recombination model? Co-estimated under a model of GC, our genome-wide average of the CO rate per-base (0.00186) is lower than the *ρ* estimate of Booker et al. (2017) (0.00924, averaged across autosomes) by a factor of 5. If we compare the total recombination rate estimated between any two adjacent bases, which corresponds to the upper bound of the recombination rate in our model (for *d* = 1 eq. 5 reduces to *ρ* = *κ* + 2*γ*), the results are quite similar (0.00841 vs. 0.00924). While this suggests that GC may contribute substantially to the *ρ* as estimated by Booker et al. (2017), this is unrealistic. LDHelmet uses long-range SNP-only data, making it attune to the broader signal of CO and less sensitive to the short-range effects of GC. Rather, the difference between the estimates likely reflects biology. The Booker et al. (2017) estimates are obtained using data from large contiguous windows of the genome aggregated over all sites and genomic partitions which vary in proportion along the genome but will be dominated by intergenic sequence. Our estimates instead reflect the (per-chromosome) recombination profile specifically for the beginning of introns. Direct recombination estimates in humans suggest that the recombination rate in introns is lower than the genome-wide average (Myers et al., 2005). Furthermore, intron length is negatively correlated with recombination rates in some taxa (Comeron and Kreitman, 2000), and our filtering strategy enriched for long introns.

### Further applications and outlook

There are several potential avenues for further work. Firstly, it should be possible to relax the assumption of an infinite sites mutation model. While our analysis of the *M*. *m*. *castaneus* data reveals very small/tolerable biases (Figure 3); basing estimates of GC and CO on more realistic mutation models might be important when analyzing more heterozygous genomes. Secondly, as a natural choice, we have assumed that loci are individual nucleotides. One could in principle extend the two locus inference to longer blocks of sequence and use the framework developed by (Lohse et al., 2011) to base inference on the joint distribution of pairwise differences. However, this comes at the cost of introducing additional assumptions and biases. Finally, it would be interesting to explore whether the machinery could be extended to three loci. If analogous analytic results for three loci are tractable, this would allow extracting substantially more signal and better estimate the rate and tract length of GC events from genomic data.

While we have limited our analyses to long introns, any genomic data partition for which pairwise heterozygosity can be accurately measured over a sufficient range of physical distances is suitable. It remains to be seen whether our method is informative about smaller genomic partitions such as centromeres and chromosomal inversions which differ from the genome-wide rates of recombination in systematic and where GC may occur but CO is restricted (Korunes and Noor, 2017).

## Acknowledgements

This work was supported by an ERC starting grant (ModelGenomLand 757648). SE is supported by an EastBio studentship from the British Biological Sciences Research Council (BBSRC). KL is supported by a fellowship from the Natural Environment Research Council (NERC, NE/L011522/1).

We thank Stuart Baird for helpful discussions and Susan Johnston for insightful comments on the manuscript.

## Supporting Information

**Figure S1:**
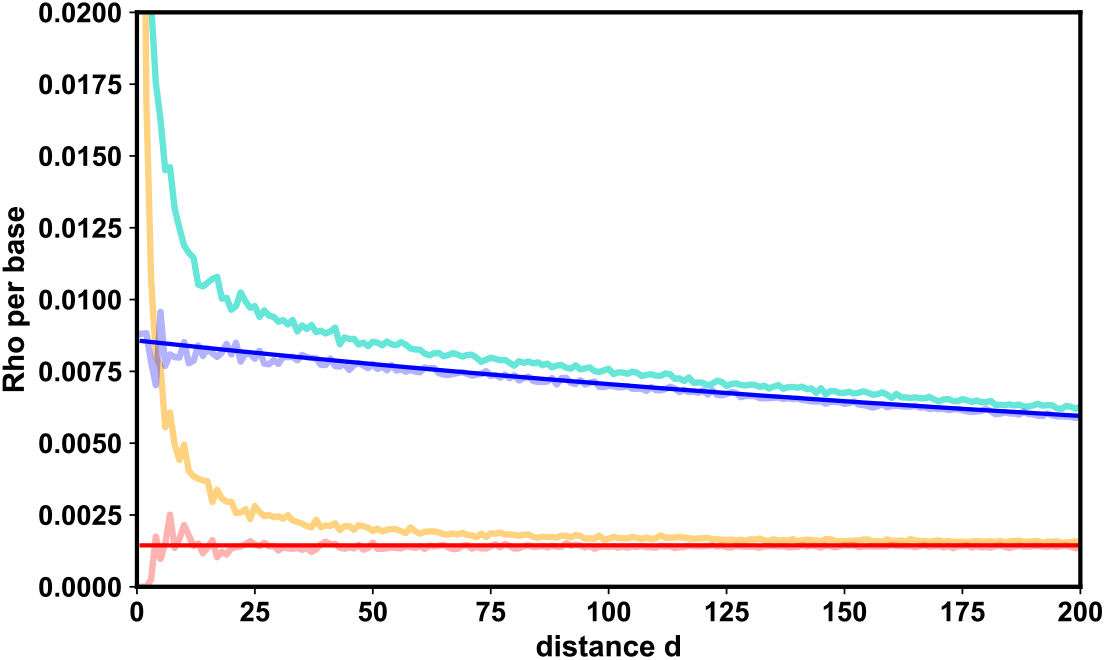
The bias of the estimator for discrete genomes. Here, we compare the analytic predictions for single-distance estimates of *ρ*/*bp* to those observed in simulations. The model predictions with and without GC are shown in dark blue and dark red, respectively. Light blue and light red show estimates from a coalescent model with a continuous genome, and the turquoise and orange lines to a discrete genome, correspondingly. Here, estimates were obtained from the combined data of 100 replicates under each simulated scenario. Parameters as in fig 1

**Figure S2:**
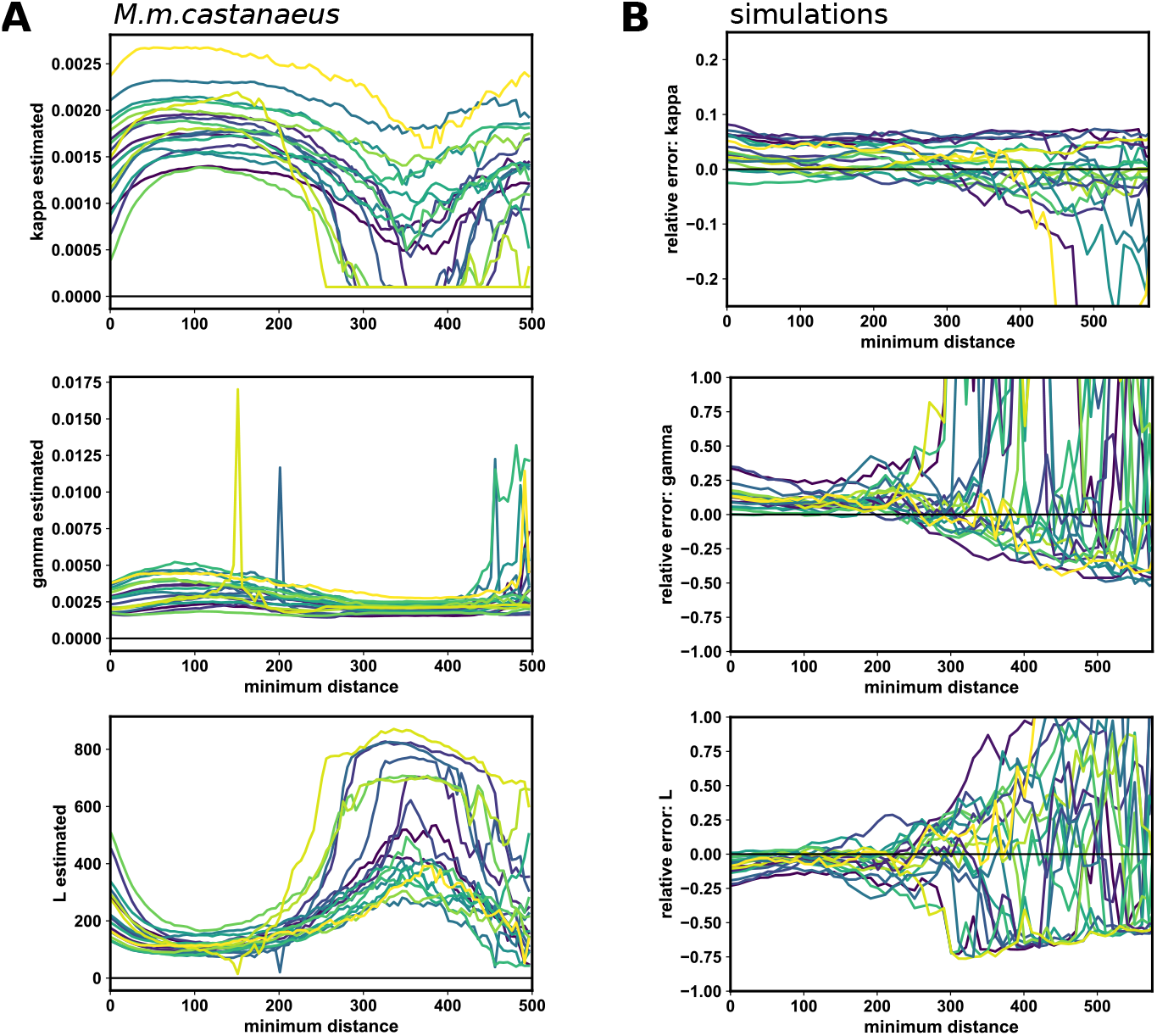
The effect of minimum distance on the per-autosome composite likelihood estimate of recombination. Panel A shows, from top to bottom, the estimated values of *γ*, *κ*, and *L* as a function of the minimum distance included in the likelihood calculation. Each indexed color corresponds to one chromosome, with chromosome 1 the darkest and chromosome 19 the lightest. Panel B shows the corresponding results obtained using the a single replicate of the simulations used for parametric boostrapping. The relative error in the estimated value, that is, the (observed - expected)/expected value, and the chromosomes are colored by index from the lowest (dark) to highest (light) recombination rate.

**Figure S3:**
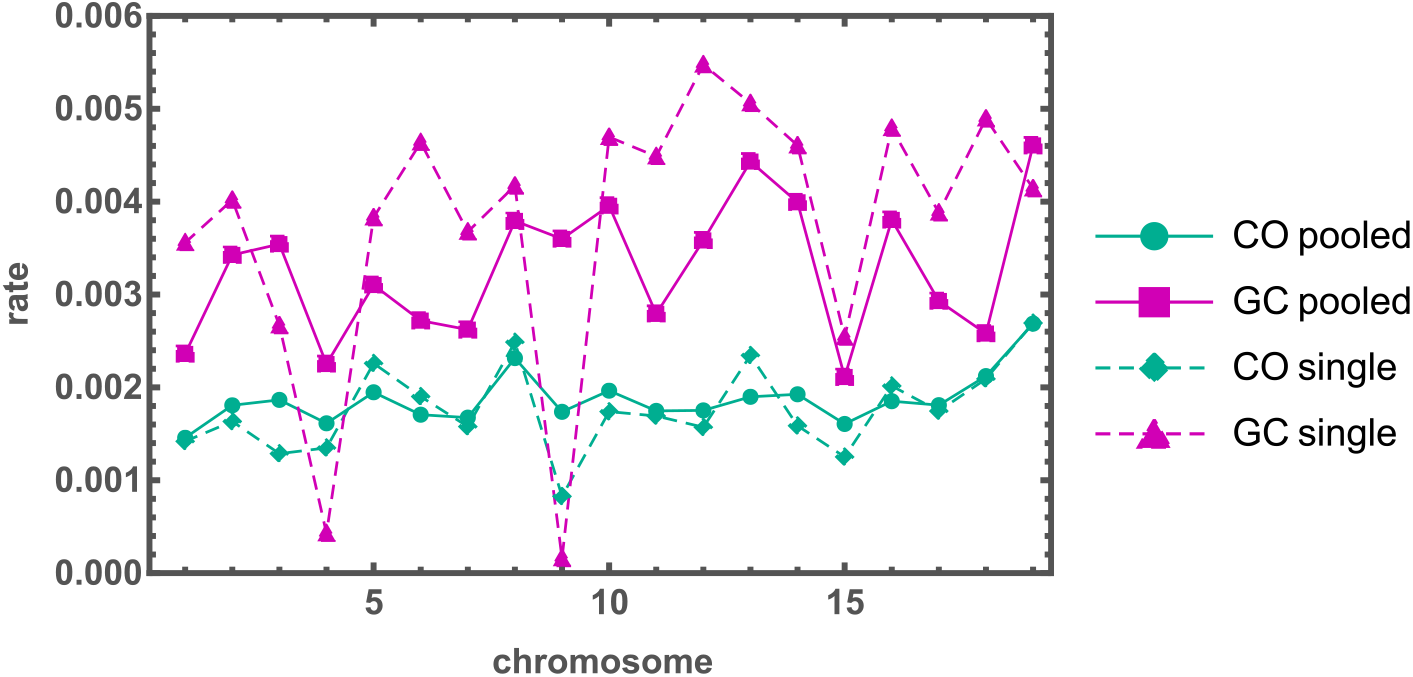
Comparison of the co-estimated CO rate (green) and GC rate (purple) *M*. *m*. *castaneus* for single-individual data (dashed) vs data pooled from ten individuals (solid). Each marker corresponds to a unique chromosome and estimate. The single- and multiple-individual estimates of the mean tract length were 107.8 and 108.4, respectively.

**Figure S4:**
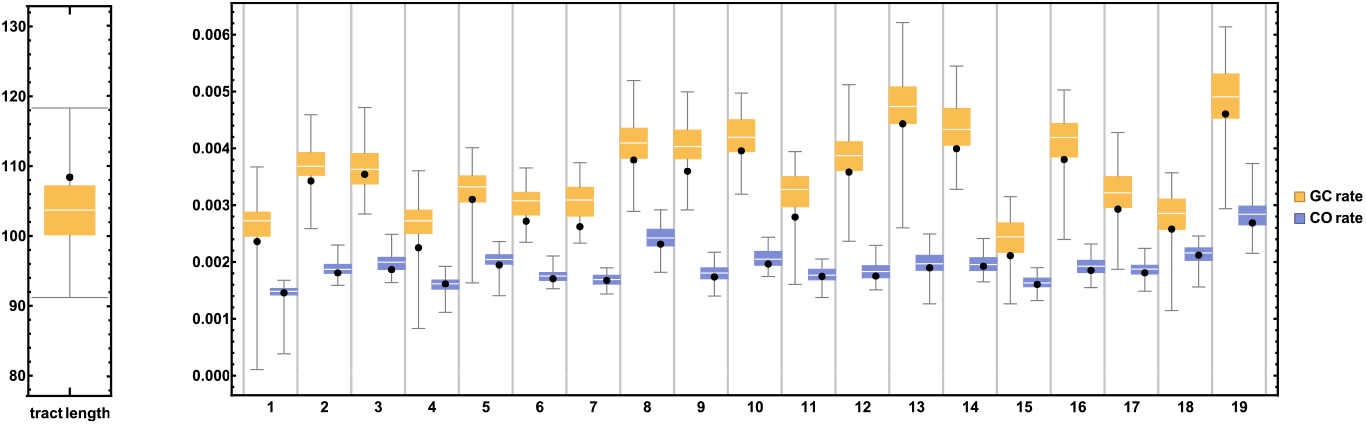
Bootstrapping results when data is limited to a single individual. The black dots correspond to the recombination parameters co-estimated for the autosomes using data pooled across ten *M*. *m*. *castanaeus* individuals (Fig. 3). Here, we randomly subsampled one individual from each of the simulation replicates. The per-chromosome GC rate and mean tract length estimates are shown in yellow, and the corresponding CO rate estimates are shown in blue.

**Figure S5:**
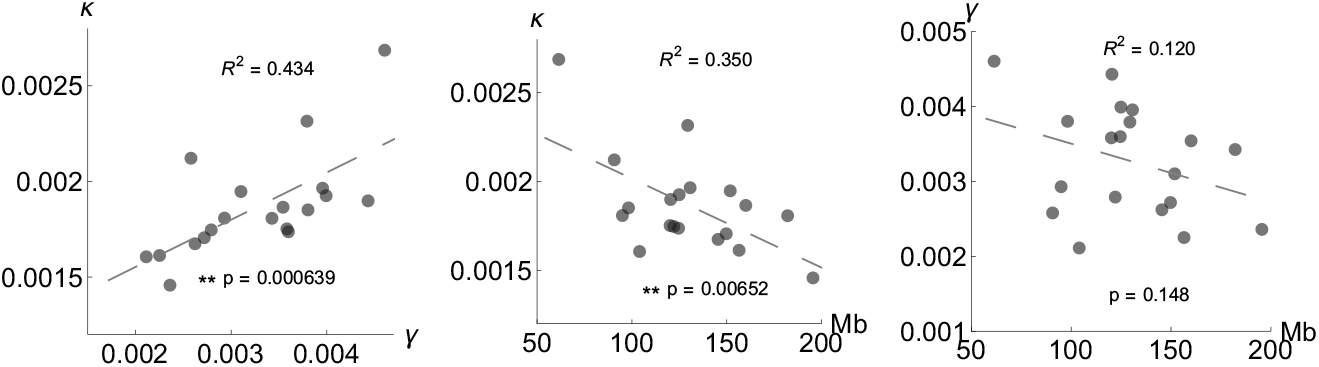
Left) Per chromosome estimates for the rates of CO and GC in *M*. *m*. *castaneus* are positively correlated; Center) Given that chromosomes have a roughly fixed map length, we expect *ρ* per base to correlate negatively with the physical length of chromosomes; Right) we find no analogous correlation between the rate of GC (*γ*) and chromosome length.

